# Evolutionary features of a prolific subtype of avian influenza A virus in European waterfowl

**DOI:** 10.1101/2021.11.24.469925

**Authors:** Michelle Wille, Conny Tolf, Neus Latorre-Margalef, Ron A. M. Fouchier, Rebecca A. Halpin, David E. Wentworth, Jayna Ragwani, Oliver G. Pybus, Björn Olsen, Jonas Waldenström

**Affiliations:** Centre for Ecology and Evolution in Microbial Model Systems, Linnaeus University, SE - 391 82 Kalmar, Sweden; Department of Virology, Erasmus Medical Centre, Rotterdam, The Netherlands; J. Craig Venter Institute, Rockville, Maryland, USA; Department of Zoology, University of Oxford, South Parks Road, Oxford, United Kingdom; Department of Pathobiology and Population Sciences, The Royal Veterinary College, AL9 7TA London, United Kingdom; Section of Infectious Diseases, Department of Medical Sciences, Uppsala University, SE751 85, Uppsala, Sweden

## Abstract

Avian influenza A virus (AIV) is ubiquitous in waterfowl, and detected annually at high prevalence in waterfowl during the Northern Hemipshere autumn. Some AIV subtypes are globally common in waterfowl, such as H3N8, H4N6, and H6N2, and are detected in the same populations at high frequency, annually. In order to investigate genetic features associated to the long-term maintenance of common subtypes in migratory ducks, we sequenced 248 H4 viruses isolated across 8 years (2002-2009) from Mallards (*Anas platyrhynchos*) sampled in southeast Sweden. Phylogenetic analyses showed that both H4 and N6 sequences fell into in three distinct lineages, structured by year of isolation. Specifically, across the eight years of the study, we observed lineage replacement, whereby a different HA lineage circulated in the population each year. Analysis of deduced amino acid sequences of the HA lineages illustrated key differences in regions of the globular head of hemagglutinin that overlap with established antigentic sites in homologous hemagglutinin H3, suggesting the possibility of antigenic differences among these HA lineages. Beyond HA, lineage replacement was common to all segments, such that novel genome constellations were detected across years. A dominant genome constellation would rapidly amplify in the duck population, followed by unlinking of gene segments as a result of reassortment within 2-3 weeks following introduction. These data help reveal the evolutionary dynamics exhibited by AIV on both annual and decadal scales in an important reservoir host.

## Introduction

Influenza A viruses have a significant impact on human and animal health worldwide. These viruses cause seasonal outbreaks and pandemics in humans, but also cause disease in domestic animals such as horses, pigs and poultry, and wild animals including birds and seals (Anthony et al, 2012; Daly et al, 2011; Vincent et al, 2014; Webster et al, 1992; Zohari et al, 2014). Central to the epidemiology of influenza A viruses are wild birds, particularly the *Anseriformes* (ducks, geese, swans) and *Charadriiformes* (shorebirds, gulls) (Olsen et al, 2006; Webster et al, 1992).Within this wild bird reservoir, influenza A viruses have been isolated from more than 105 species across 26 different families. Wild birds maintain a large diversity of avian influenza A viruses (AIV) subtypes; 16 of the possible 18 haemagglutinin (HA) subtypes and 9 of the 11 neuraminidase (NA) subtypes are found in wild birds (Olsen et al, 2006; Olson et al, 2014). Of importance in the context of emerging and re-emerging disease is that this large diversity of AIV in waterfowl frequently spillover into poultry (Verhagen et al, 2017), which may facilitate the introduction of AIVs into future human pandemic viruses (Smith et al, 2009; Wille & Holmes, 2020).

In the Northern Hemipshere, waterfowl have high AIV prevalence in the autumn, linked to the congregation of migrating birds including large numbers of immunologically-naïve juveniles (Latorre-Margalef et al, 2014; van Dijk et al, 2014). A large diversity of HA subtypes are maintained in waterfowl, however certain HA-NA subtype combinations, such as H3N8, H4N6, H6N2, are over-represented at waterfowl surveillance sites and may comprise the majority of viruses detected and isolated. Conversely, a number of HA-NA subtype combinations are uncommon or absent from these study sites (Latorre-Margalef et al, 2014; Munster & Fouchier, 2009; Wilcox et al, 2011; Wille et al, 2018). The consistent detection of diverse subtypes, combined with low rates of evolutionary change of amino acid sequences led early studies to postulate that avian AIV was in evolutionary stasis (Hatchette et al, 2004; Sharp et al, 1997; Webster et al, 1992). However, high rates of molecular evolution (~10^−3^ substitutions per site per year) and inference of positive selection (dN/dS) in AIV genomes have firmly refuted this hypothesis (Chen & Holmes, 2006).

In the last decade, a number of interconnected hypotheses of AIV evolution have been put forward based on observed AIV genetic structure. Central to these hypotheses are the evolutionary features of antigenic drift and antigenic shift. Antigenic drift is the fixation, by natural selection, of mutations in the HA and NA that enable the virus to evade the host immune response. These mutations arise through an error prone RNA-dependant RNA polymerase (Chen & Holmes, 2006; Gething et al, 1980). Antigenic shift, or process of reassortment following coinfection allows for the generation of novel genome constellations with altered antigenic properties (Lowen, 2017; Steel & Lowen, 2014). Together, these processes result in the capacity for rapid genetic and antigenic change, and while the same subtypes may appear annually in a population, these subtypes consist of multiple genetic lineages, which accumulate mutations, resulting in high levels of genetic diversity in the viral population over time.

In this study, we aimed to reveal the genetic and evolutionary features that allow for the long-term maintenance of AIV subtypes in waterfowl popuations. We collected and sequenced 248 H4 isolates across 8 years from migratory Mallards (*Anas platyrhynchos*) at a stopover site in SE Sweden. H4 is not only the most common AIV in European ducks (Latorre-Margalef et al, 2014; Munster et al, 2007), but also in Asian wild and domestic ducks (Cheng et al, 2010; Kang et al, 2013; Wisedchanwet et al, 2011; Zhang et al, 2012). We used Mallards as a model, as they are considered to be an important reservoir for AIV diversity and have formed the basis of the current understanding of spatial and temporal avian AIV dynamics in nature (Latorre-Margalef et al, 2014; Olsen et al, 2006). Specifically, from these data, we aimed to reveal the role of different genetic lineages, examine how this genetic variation may relate to antigenic differences, and address the role of reassortment in both within-year and among-year maintenance of this virus subtype. We suggest that a complex combination of natural selection, genetic hitchhiking, and reassortment is responsible for the evolutionary patterns of AIV observed in wild birds.

## Methods

### Ethics Statement

All trapping and handling of mallards was done in accordance with regulations provided by the Swedish Board of Agriculture under permits from the Linköping Animal Research Ethics Board (permit numbers 8-06, 34-06, 80-07, 111-11, and 112-11).

### Study Site and Virus Collection

Wild Mallards were captured as part of an ongoing long-term AIV surveillance scheme (Latorre-Margalef et al, 2014) at Ottenby Bird Observatory, Sweden (56° 12’N, 16° 24’E). Details on trapping, AIV surveillance and virus isolation have been published previously (Latorre-Margalef et al, 2016; Latorre-Margalef et al, 2014). Briefly, for each captured duck, either a cloacal swab or a faecal sample was collected and placed in virus transport medium (VTM). All samples were stored in a −80 °C freezer within 2-6 hours of collection. Viral RNA was extracted from the VTM samples, and assayed for a short fragment of the matrix gene using reverse-transcription real-time PCR (rRT-PCR) (Spackman et al, 2002). Samples positive for AIV were inoculated into 10-12 day old specific pathogen free (SPF) embroynated chicken eggs by the allantoic route. HA subtypes of positive samples were determined using hemagglutination inhibition, and NA subtypes were determined by sequencing (Latorre-Margalef et al, 2016; Latorre-Margalef et al, 2014).

### Sequence Dataset

Full genomes were sequenced as part of the Influenza Genome Project (http://gcid.jcvi.org/projects/gsc/influenza/index.php), an initiative by the National Institute of Allergies and Infectious Diseases (NIAID) as previously described (Wille et al, 2018). Briefly, AIV RNA was extracted and the entire genome was amplified using a multi-segment RT-PCR strategy (Zhou et al, 2009; Zhou & Wentworth, 2012). The amplicons were sequenced using the Ion Torrent PGM (Thermo Fisher Scientific, Waltham, MA, USA) and/or the Illumina MiSeq v2 (Illumina, Inc., San Diego, CA, USA) instruments. When sequencing data from both platforms was available, the data were merged and assembled together; the resulting consensus sequences were supported by reads from both technologies.

We attemped sequencing 289 viruses, and 248 were successfully sequenced. Viruses that were not successfully sequenced did not pass QC at the sequencing facility, and we did not attempt resubmission of samples or resequencing.

All sequences generated in this study have been deposited in GenBank (accession numbers CY164135-CY166103).

### Phylogenetic Analysis

Sequences generated in this study, in addition to Eurasian reference sequences mined from the Influenza Research Database (http://www.fludb.org/) were aligned using MAFFT (Katoh et al, 2009). Maximum likelihood trees were used to explore the temporal signal and clock-like behaviour of each data set by performing linear regressions of root-to- tip distances against year of sampling, using TempEst (Rambaut et al, 2016). Using BEAST v10.4 or v1.8, time-stamped data were analysed under the uncorrelated lognormal relaxed molecular clock (Li & Drummond, 2012), and the SRD06 codon-structured nucleotide substitution model (Shapiro et al, 2006) and the Bayesian skyline coalescent tree prior. Three independent analyses of 100 million states were performed, which were then combined in LogCombiner v1.8 following the removal of a burnin of 10 per cent. Convergence was assessed using Tracer v1.6 (http://tree.bio.ed.ac.uk/software/tracer/). Maximum credibility phylogenies were generated using TreeAnnotator v1.8 and visualized in FigTree v1.4. The N2 phylogeny was estimated using Mr Bayes 3.2.1 (Ronquist et al, 2012). All trees were run until convergance, which was assessed using Tracer v1.4. The tree was visualised using FigTree v1.4.

Phylogenetic lineages were identified through a combination of pairwise identity plots (summarized in Fig S1) and phylogenetic tree shape. From these analyses, 95% sequencing identity was established as a cut-off for genetic lineage delineation, similar to previous assessments (Huang et al, 2014; Reeves et al, 2011; Wille et al, 2013). Lineage proportion plots were plotted using *ggplot2* library in R v. 3.5.1 integrated into RStudio v. 1.0.143 .

Discrete trait analysis was performed using the asymmetric trait evolution model, and social networks were inferred with Bayesian Stochastic Search Variable Selection (BSSVS) (Lemey et al, 2009). Connectivity among locations was determined using Bayes Factor analysis as implemented in SpreaD3 (Bielejec et al, 2016). We considered Bayes Factors of greater 10 to be strong support, and greater than 100 to be decisive support (Hill et al, 2021; Jeffreys, 1961).

Initial computations were performed on resources provided by SNIC through Uppsala Multidisciplinary Center for Advanced Computational Science (UPPMAX) under project b2013122.

### Comparison of H4 HA1 peptide sequences of Swedish H4N6 virus

In order to evaluate possible antigenic properties governing between different lineages of H4, HA amino acid sequences of each lineage were aligned using MAFFT, and consensus sequence for each lineage determined using Geneious Prime 2020.2.4. Consensus sequences from different years were aligned to identify substitutions that had been introduced between years. In these alignments, sequence regions corresponding to established antigenic sites (A to E) in the closely related and well studied H3 HA1 was indicated as a reference (Broecker et al, 2018). These substitutions were also visualized in the HA monomer derived from the crystal structure of an AIV H4 HA (PDB ID: 5XL1), using Geneious Prime 2020.2.4.

## Results

### H4 appears annually, at high frequency, in migratory Mallards

As part of a long-term AIV surveillance scheme, 18,643 cloacal/fecal samples from migratory Mallards were collected between 2002–2009 at Ottenby Bird Observatory in southeast Sweden, as outlined in (Latorre-Margalef et al, 2014). From these samples, 289 H4 viruses were successfully isolated, representing 27% of all viruses isolated from Mallards in the study period (Fig 1).

**Figure 1.**
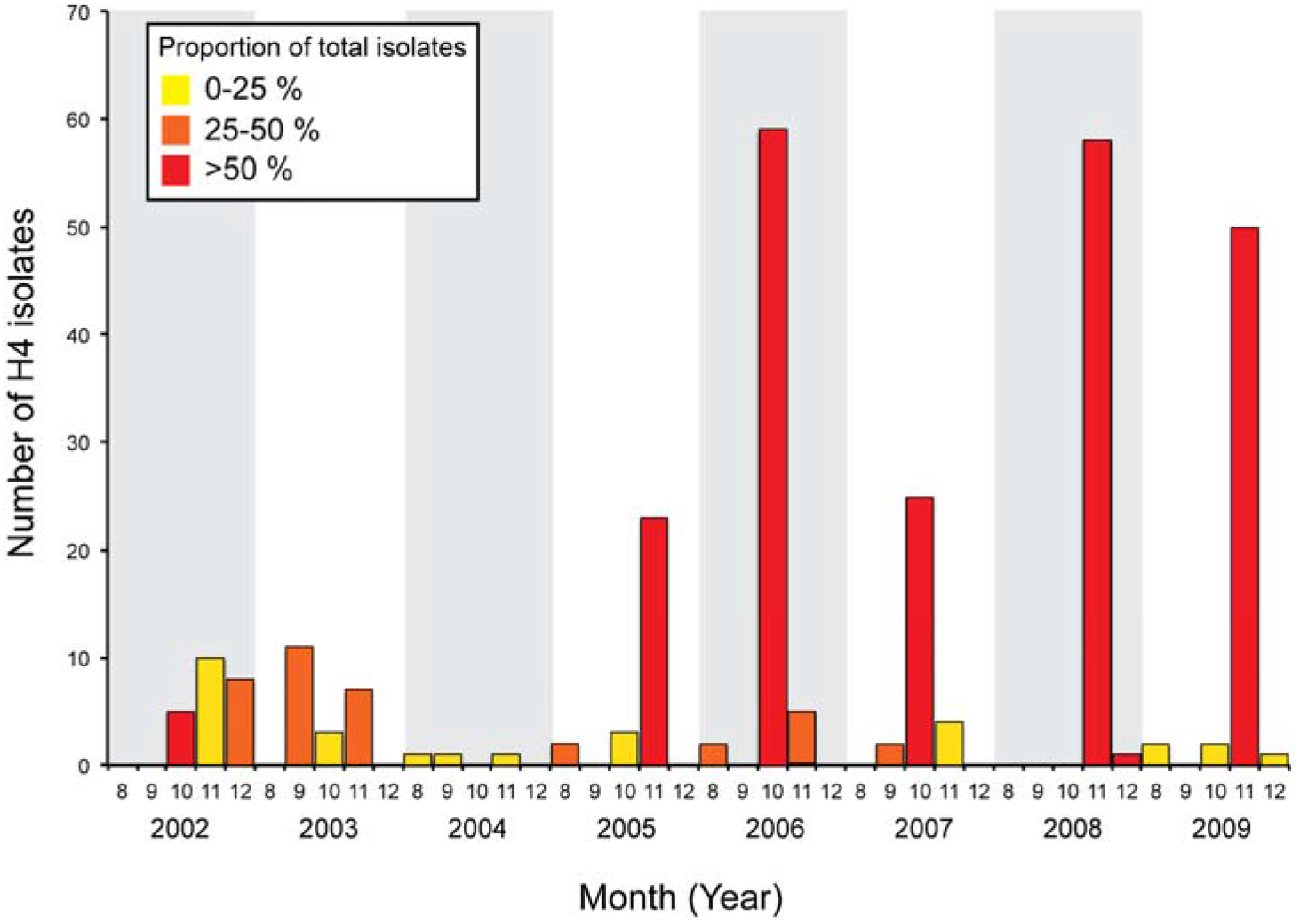
H4 AIVs are isolated at high frequency and constitute a substantial proportion of viruses from Mallards sampled during the autumn migration at Ottenby Bird Observatory.

For 2005–2009, the years for which we have the most data, we find “outbreak-like” patterns of H4 in the population, wherein during October or November, this subtype comprised > 50% of all isolated viruses (Fig 1). In 2002 and 2003, years in which we have less data, we find a bimodal H4 prevalence peak, with high prevalence in September and November, respectively. No prevalence pattern could be observed in 2004 due to limited number of isolates (Fig 1). Moreover, for H4 viruses isolated from 2002 to 2009, all NA subtypes except N4 and N7 were detected. The majority (79%), however, were H4N6 viruses (Fig S2A).

### Long term phylodynamics of H4N6

A total of 246 H4 viruses isolated from 2002–2009, were sucessfully sequenced. Reconstruction of the H4 phylogenly of Eurasian sequences revealed two main lineages: one lineage comprising sequences from Asia and Australia (1980–present) (Fig 2A, collapsed node) and a second lineage dominated by European sequences, most of which were generated in this study, in addition to a smaller lineage of Asian sequences (Fig 2A). The estimated time of the most recent common ancestor (tMRCA) of all Eurasian sequences, including both the Asian/Australian and European lineages, is 1926–1957 (95% higest posterior density [HPD], mean 1946).

**Figure 2:**
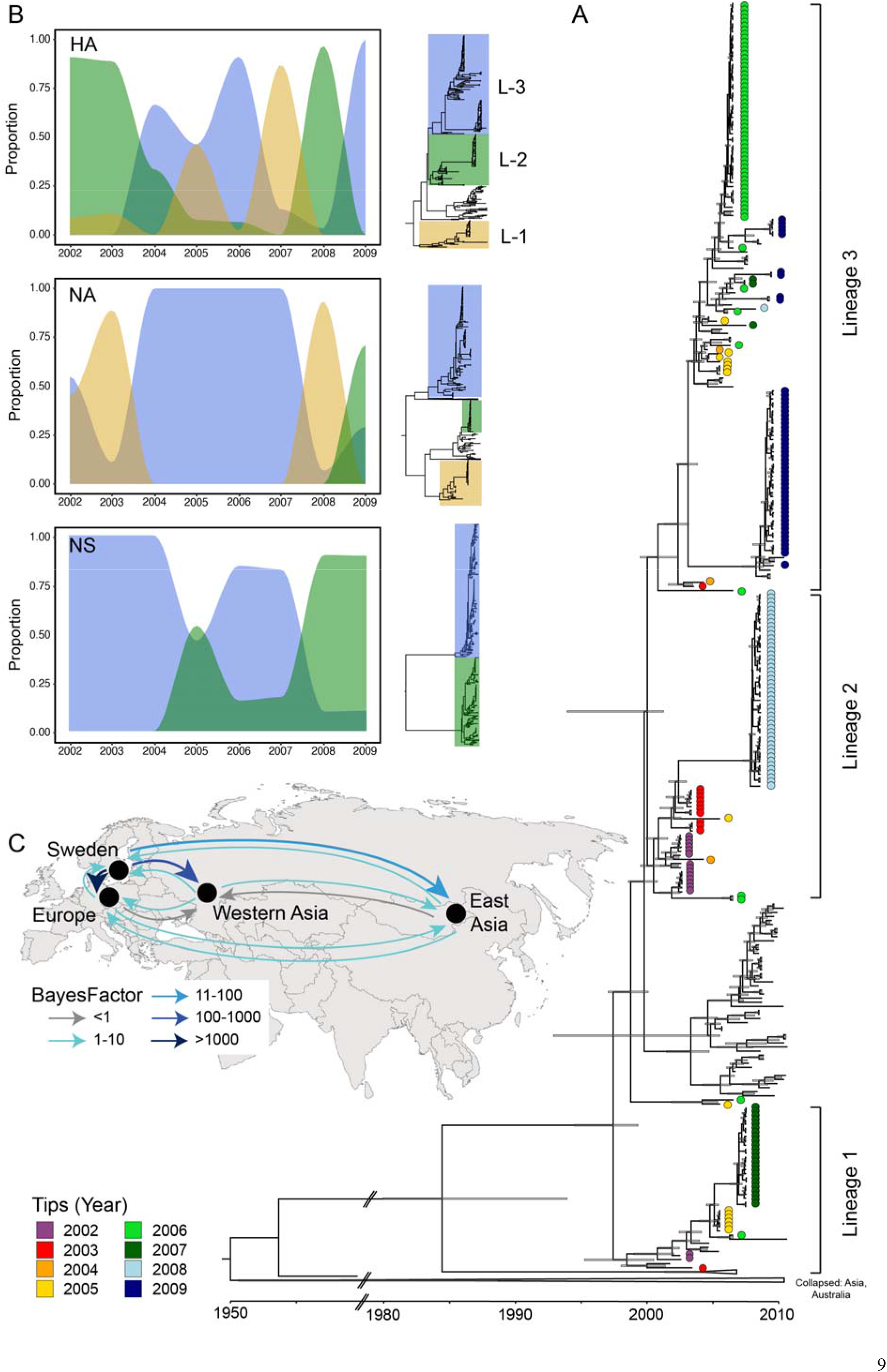
Features of long-term evolution of H4. (A) Time-scaled phylogeny of the Eurasian lineage of H4. Phylogeny includes all H4 sequences in GenBank from 1950-2010. Sequences generated in this study are denoted by a filled circle, with different colours referring the year of sample collection. Branch nodes correspond to the 95% HPD of node height. (B) Proportion of lineages of each H4, N6, and NS detected in Sweden between 2002 and 2010. For each segment the phylogeny with highlighted lineages is shown. For NS, the lineages correspond to the A and B alleles. In both (A) and (B), lineages circulating in Sweden are refered to as Lineage (L) 1-3 for clarity only. (C) Viral diffusion between geographic locations. Sequences included in Europe include Netherlands, Czech Republic, and England; Western Asia includes western Russia and the Republic of Georgia; Eastern Asia includes China, Japan, Korea, Mongolia, Russian Siberia. Arrow colour refers to strength of support for inclusion in a phylogeograhic model (*i.e.* Bayes Factors) where values greater than 10 represent strong support, whereas values greater than 100 are considered decisive support.

Given that two-thirds of H4 sequences from Eurasia between 2000–2009 are sequences generated in this study, the European lineage of the H4 phylogenetic tree is largely defined by the sequences from our study site. In addition to these, there are a smaller number of sequences from the Netherlands and Czech Republic in GenBank. All H4 sequences from Sweden are within 95% sequence identity (Fig S1) and the European lineages comprising the sequences generated in this study diverged at 1997.7 (tMRCA, 95% HPD 1994–1999) (Fig 2A). We find the sequences generated in this study to fall into 3 lineages with the exception of four sequences (including two sequences from 2010) that group closely with Asian sequences nested within this European lineage. We refer to those lineages detected in Swedish Mallards as Lineage 1–3 for clarity only (Fig 2A).

There are three general observations regarding the distribution of sequences across the three lineages. First, sequences from 2002 and 2003 fall into the same lineage (Lineage 2). Second, unlike all other years in this dataset, sequences from 2005 fall into two disctinct lineages with equal proportion: the lineage comprising sequences from 2006 (Lineage 3) and that of 2007 (Lineage 1). And finally, starting in 2006 (for which we have the most data), sequences from within each year fall into a single lineage with ~99% nucleotide similarity, and that these lineages are distinct from the dominating lineage in the prior and subsequent year. Indeed, the lineage containing sequences in 2006 and 2009 (Lineage 3), do not contain more than 3 sequences from intermediate years. Taken together, we find that lineage replacement in H4 is high between years (Fig 2B).

To attempt to better elucidate the source of annual viral introduction into Sweden, we undertook a phylogeographic analysis. Based on the available sequences from 1980-2010, there is strong evidence for virus movement between Sweden and other European locations (Bayes factor 88067.9, posterior probability 1), between Sweden and western Asia (here western Russia [Moscow] and Republic of Georgia) (Bayes factor 264, posterior probability 0.987), and between Sweden and eastern parts of Asia (Bayes factor 24.7, posterior probability 0.88). In this analysis, virus movement outside Sweden was poorly supported (Bayes factor <10 and posterior probability <0.5), most likely because of the low number of sequences from other regions (Fig 2C). Overall, due to poor sequence coverage from Eurasian locations outside of Sweden it is challenging to infer the role of different geographic locations in the introduction of viruses into Swedish-caught Mallards.

Similar to the H4 tree, sequences from this study play an important role in defining the shape of the Eurasian N6 phylogenetic tree (Fig S2). The N6 lineages are more distantly related, as compared to H4 lineages, where the three distinct N6 lineages detected in Ottenby are ~93% identical and postulated to have evolved from their MRCA ranging from 1977–1987 (95% HPD). The second most common NA subtype identified was N2, and as with N6, Swedish N2 sequences were similar to those circulating in Europe, most often Sweden, during the same time period. N2 sequences from 2005 and 2008 formed discrete lineages sharing 99% sequence identity, whereas sequence variation was greater in 2002 and 2007 (Fig S3). In addition to N6 and N2, the dataset included N1 (n=1), N3 (n=3), N5 (n=1), and N9 (n=3). In all instances these sequences were most closely related to other European AIV sequences, and the top 10 Blast matches often included other sequences from Sweden (Table S1).

Globally, non-structural protein (NS) sequences fall into two major lineages, termed allele A (including viruses with broad host range) and allele B (viruses specific to birds). NS sequences of viruses sequenced in this study fell into both allele A and B, and were most similar to NS sequences of AIV from Eurasian wild birds (Fig S4). Startgin in 2015, both alleles were found in the population each year, although with differing proportions. The shapes of the phylogenetic trees for the “internal” segments (PB2, PB1, PA, NP, M) were similar to those previously characterized (Fig S5). Within the “internal” segments, we found a pattern similar to that of HA and NA, wherein sequences from 2006-2009 fell into discrete year-specific lineages.

### Structural variation in HA protein sequences of Swedish H4N6 virus

To evaluate putative antigenic differences in amino acid consensus sequences of these lineages, genetic changes were evaluated by aligning H4 HA sequences across years and by mapping identified substitutions onto the crystal structure of H4 (Fig 3). In line with the nucleotide phylogeny, the phylogeny of HA1 amino acid consensus sequences from 2006–2009 exhibit temporally discrete lineages. Across years, amino acid sequence variation predominately occured in the HA1, and more specifically in, or in close proximity to, regions corresponding to antigenic sites identified in the H3 HA1 (Broecker et al, 2018). While antigenic sites for H4 have not been mapped, H3 and H4 belong to the same lineage within group 2 HA subtypes (Latorre-Margalef et al, 2013), and we therefore postulate that these sites should be rather conserved across these two subtypes.

**Figure 3:**
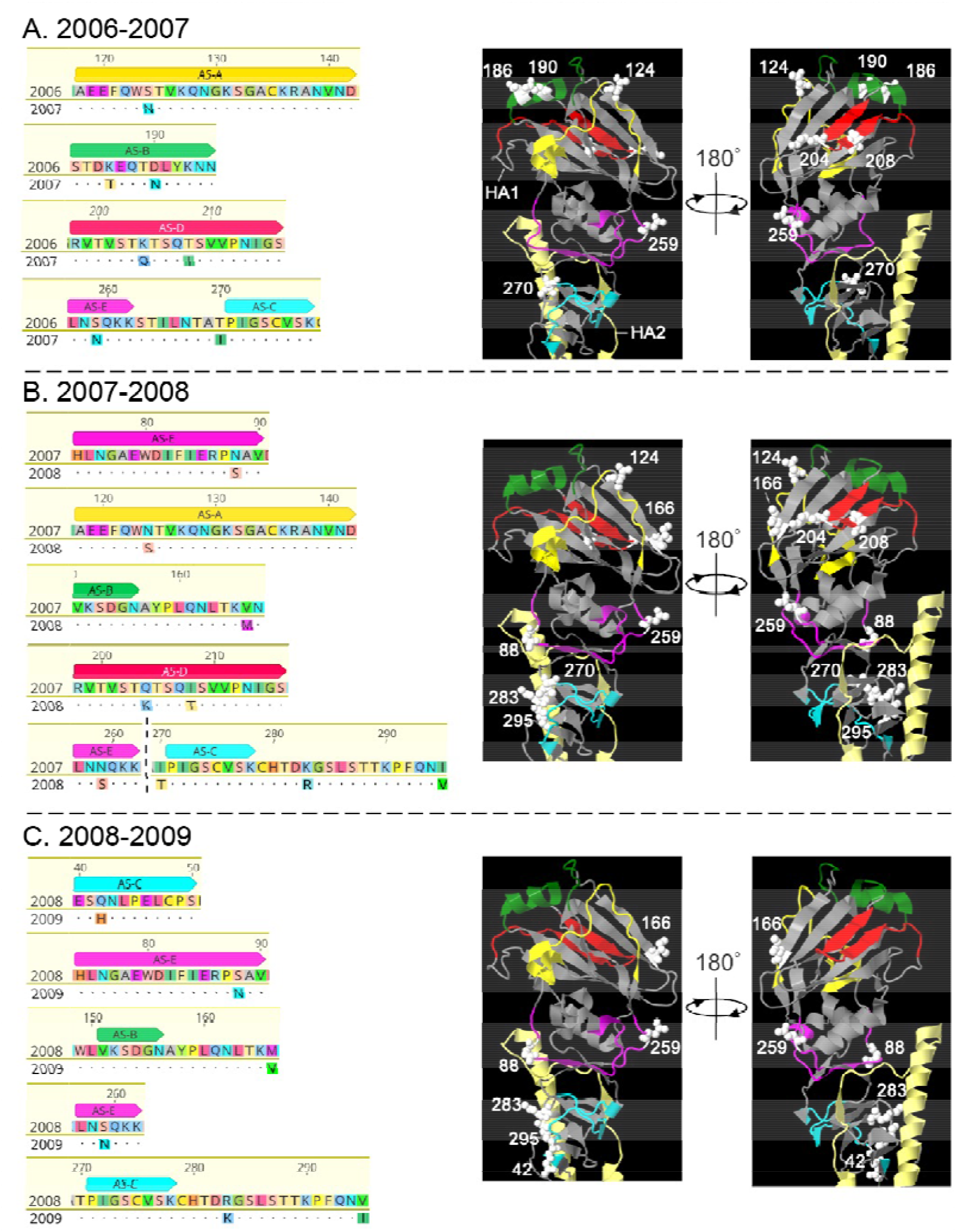
Amio acid substitutions across years between distinct H4 HA1 lineages of Swedish H4N6 strains. Specfically, amino acid substitions between sequences from (A) 2006-2007, (B) 2007-2008, and (C) 2008-2009. Amino acid sequence alignments showing substitutions between years indicated in, or in close proximity to, regions corresponding to established antigenic site A-E (AS-A – AS-E) in the HA1 peptide of H3 (Broecker et al 2018). In sequence comparisons, consensus sequences from 2006, 2007 and 2008 serve as reference and substitutions in sequences from the following year is indicated by single letter abbreviation of amino acids whereas identical residues are represented by dots. In B, the dotted vertical line indicates a truncation for clarity only. Substitutions visualized by white colour in the crystal structure of H4 HA1. Antigenic site regions are indicated and colour coordinated with corresponding regions in alignments. HA1 (grey) and HA2 (light yellow) peptide monomers are indicated the left structure of the 2006-2007 panel

When assessing the amino acid changes between consecutive years, we identified between 6–9 amino acid substitutions in the HA1 region. Specifically, between 2006–2007 (Lineage 3-> Lineage 1, Fig 3A) we found seven substitutions, 2007–2008 (Lineage 1-> Lineage 2, Fig 3B) nine substitutions and 2008–2009 (Lineage 2-> Lineage 3, Fig 3C) six substitutions. When comparing the sequences from 2006 compared to 2007, amino acid substitutions were located around the top of of the globular head of HA1, close to secondary structures forming the receptor binding site, including the 130 loop, the 190 helix and the 220 loop, which partly overlap with H3 antigenic site A, B and D, respectively (Fig 3A). The majority of these substitutions are functionally favourable (Table S2), with the exception of K186T and T208I. In the transition from 2007 to 2008, four out of nine substitutions are found in loop structures closer to the stem region of HA1, including the 260 loop. These include Asn88Ser, Ser259Asn in a region corresponding to antigenic site E in H3 as well as I270T, K283R and I295V adjacent to site C (Fig 3B). All substitutions except I208T and I270T, are functionally favourable. It is noteworthy that four substitutions in position 124, 204, 208, 259 and 270 are reversions to amino acid residues expressed in the 2006 consensus sequence. Finally, in the transition between 2008 and 2009, the only unique substitution is Q42H located in what is consistent with antigenic site C in H3 HA1. This substitution is neutral from a functional standpoint. In addition, five of six substitutions found in the consensus sequence from 2009 compared to that from 2008 represent reversions to amio acids expressed in the 2007 sequence. These include amino acid at position 88, 166, 259, 283 and 295 (Fig 3C). For clarity, antigenic sites A-E correspond to those in the HA1 peptide of H3 as there is no characterisation of H4 available (Broecker et al, 2018).

### Lineage replacement and no perpetuation of genome constellations across years

Analysis of temporal changes of H4N6 genome constellations revealed frequent lineage replacement across years (Fig 2B). Plots of lineage proportions of the glycoproteins and the NS alleles illustrate that, while lineage replacement is frequent for HA, concomitant replacement among multiple segements occurred less frequently (Fig 2B). For example, between 2003–2004, and 2008–2009, there was a replacement of both HA and NA lineages, and between 2007–2008 there was a simultaneous replacement of HA, NA and NS (Fig 2B). The replacement between 2007 and 2008 was not limited to only segments with established antigenic properites (HA and NA), but rather new lineages were introduced for all segments in addition to the replacement from NS allele A to allele B (Fig 2B, Fig S2, S4-S5). This global replacement also resulted in a reintroduction of lineages in 2008 that were circulating among sampled birds six years earlier, in 2003.

Bringing all lineage data together, we may assess whether genome constellations remain in the population across multiple years (Fig 4, Fig S6). Importantly, despite not detecting annual global replacements, there was sufficient lineage replacement across the different segments for novel constellations to proliferate each year. That is, genome constellations identified in one year, were not found again in the population the following year (Fig 4, Fig S6). Using 2005–2006 as an example: no constellations identified in 2005 were detected again in 2006, despite the fact that some H4 sequences from 2005 were found in the same phylogenetic lineage as sequences from 2006. In this case, the lineages of four gene segments were different between the dominant genome constellation in 2005 compared to the dominant constellation from 2006 (Fig 4). This example is not an outlier, rather is consistent with findings between all years of this dataset (Fig S6)

**Figure 4:**
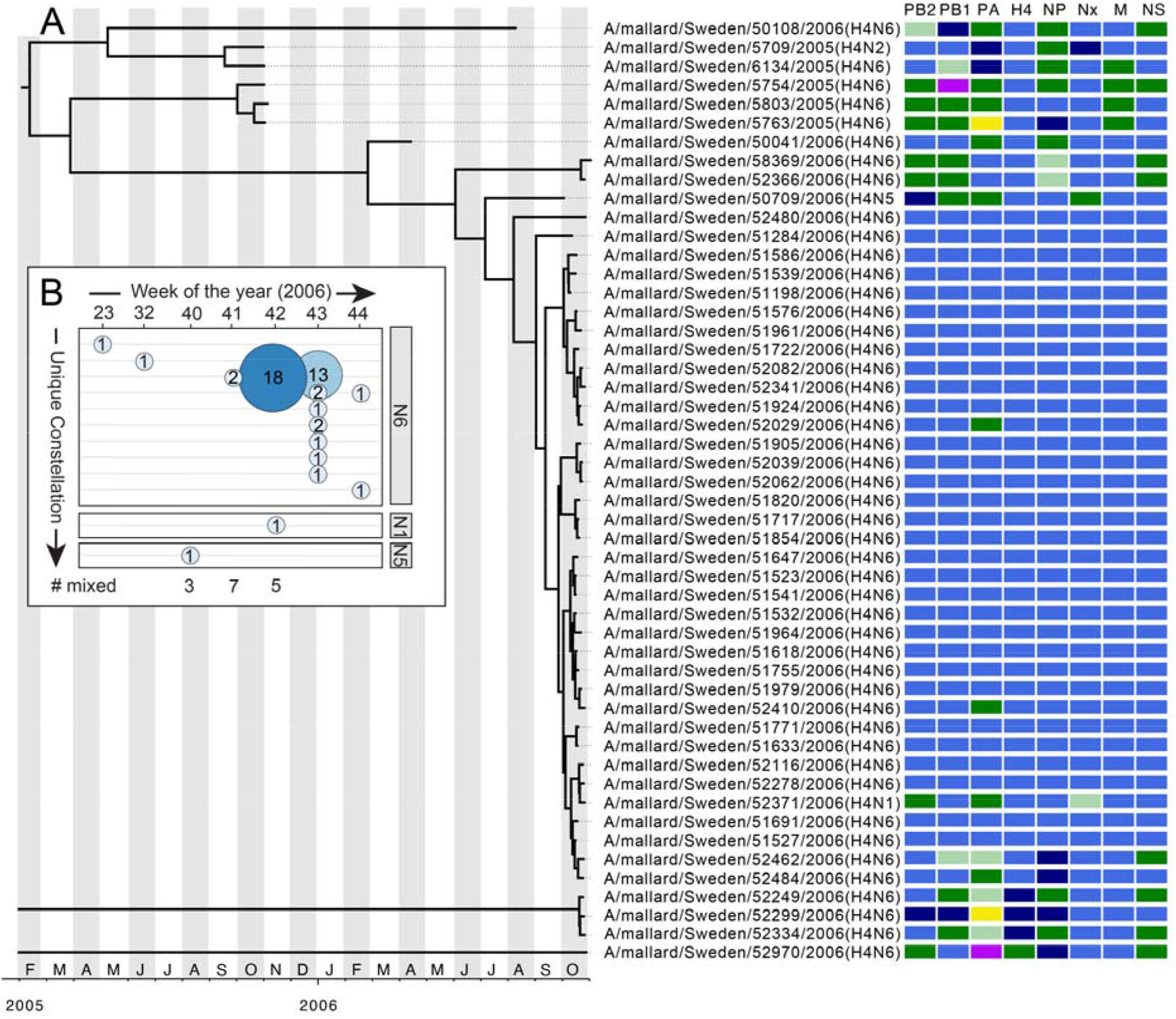
Within-year reassortment. (A) Time-structured phylogeny of H4 sequences from 2005 and 2006 comprising Lineage 1. Scale bar indicates time, with year and month indicated. Four H4 sequences from viruses detected in 2006 did not fall into this lineage, and therefore tMRCA is not shown in order to preserve resolution. Each tile column represents a segment, ordered by size: PB2, PB1, PA, H4, NP, Nx, M and NS, and each row comprises the genome constellation of a virus. Different colours pertain to different viral lineages. Genome constellations for all viruses is presented in Fig S6. (B) Genome constellation for each week of 2006. Week number is presented on the X axis and each unique constellation is on the Y axis. Bubble size and colour refer to the number of times each genome constellation was detected. Different NA subtypes are in separate panels. At the bottom the number of mixed H4 infections detected in each week is indicated. Change of genome constellations over time for all years is presented in Fig S7.

### Rapid unlinking of genome constellations within year

Within each year, particularly between 2006–2009, we identified a number of different genome constellations. Despite the diversity, more than 50% of viruses characterized within each year comprised a single, dominant constellation (Table S3). Data from 2006–2009 further indicate that the first genome constellations detected each year were not the most frequently-observed constellation overall that year. Following the introduction of the dominant constellation into the population, it was consistently detected and isolated from Mallard samples for 1-2 weeks, followed by a period (1–2 weeks) wherein this constellation rapidly unlinked and a diversity of new genome constellations were detected in the population. These novel and diverse constellations most likely contained gene lineages originating from other cocirculating viruses with different HA-NA subtype combinations and formed through the process of reassortment (Fig 4, Fig S6-S7).

Using 2006 as an example, the first virus constellations were detected in week 23 followed by a detection in 32. These constellations were not detected again, and were replaced by a frequently-detected constellation that accounted for 73% of isolates (Fig 4A) in week 41. This constellation initially appeared at low numbers in week 41 (n=2 sequenced isolates), however it was the only H4 genome constellation detected in week 42 (n=18 sequenced H4 isolates) and remained at high proportion relative to other constellations in week 43 (n=13 sequenced H4 isolates). Seven other genome constellations were also identified in week 43, and one additional novel constellation was detected in week 44 (Fig 4). Data from 2005 revealed a genetic variaton that was different compared to other years, wherein isolates comprised high levels of constellation diversity, presumably due to the presence of two different HA lineages and NS alleles (Table 1, Fig S6). In 2006, we detected 15 “mixed” viruses – these are viruses containing more than one sequence for any one segment. These mixed infections are likely reflective of co-infection in the Mallard host, but as we sequenced viruses propagated in eggs we cannot make strong inferences about these viruses (Fig 4, Table S4). These mixed viruses spanned the periods of highestprevalence and HA/NA diversity during the autumn.The year 2006 accounted for a large proportion of mixed viruses in our dataset, and in this year there was substantial cocirculation of other viral subtypes in the Ottenby duck population (*e.g.* H1N1, H3N6, H6N2; Latorre-Margalef *et al* 2014) (Fig 4, Table S4)

## Discussion

### How are subtypes maintained in waterfowl on a decadal scale ?

Low pathogenic AIV H4N6 is one of the most abundant virus subtypes isolated from waterfowl in Europe. Indeed, these viruses have been circulating at high frequency in migratory Mallards at our study site, in southeast Sweden, since the start of AIV surveillance in 2002, and continue to circulate at high frequency today (Latorre-Margalef et al, 2014; Munster et al, 2007; Wallensten et al, 2007). An important question, however, is how this subtype is maintained in a Mallard population across many years, given that high rates of infection should confer immunity in the population. Through the analysis of a dataset comprising viruses from the same species and location, but across multiple years, we were able to reveal key genetic features likely playing a key role in this phenomenon.

Unlike human influenza A virus, AIV HA segments do not exhibit a classic “ladder-shaped” phylogeny (Fitch et al, 1997). These ladder-shaped phylogenies are the result of strong positive selection on the HA, leading to antigenic drift, or the fixation of mutations in the HA (and NA) that enables the virus to evade the immune system (Koel et al, 2013; Vijaykrishna et al, 2015; Wille & Holmes, 2020). Rather, multiple lineages co-circulate in wild bird populations (Chen & Holmes, 2006, 2010). Herein, we detect 3 lineages of H4 in the mallard population. These lineages are genetically similar (within 95% nucleotide similarty), although none are present in all years. We find a phenomenon whereby there is a different HA lineage circulating each year, and within years 2006-2009, one lineage comprises ~90% of all isolates sequenced. Indeed, this is reflective of antagonistic co-evolution in a sympatric model of circulating AIV, wherein host immune pressure will drive phylogenetic branching of viruses into discrete antigenic lineages, and eventually, subtypes, with limited overlap in antigenic space (Recker et al, 2007). Alternatively, or in concert, these patterns could rather be driven by migration mediated metapopulation dynamics, which frequently re-introduces lineages to the site.

Our data is consistent with the hypothesis proposed by Chen & Holmes (2010), whereby genetic structure of AIV is shaped by a combination of occasional selective sweeps in the HA and NA segments, coupled with transient genetic linkage to the internal gene segments. Indeed, we identified annual replacement of HA, although simultaneous replacement of HA, NA and NS were less frequent. While NS is less studied as a target for cellular and humoral immune responses, there are several reports on the inhibitory effect of NS1 on the innate immunity and fitness differences of the different alleles (Adams et al, 2013; Gack et al, 2009; Nogales et al, 2018; Rajsbaum et al, 2012; Yodsheewan et al, 2013). In 2008, we detected lineage replacement in all segments, which coincided with the shift of NS A to B allele in the virus population. This event does support the transient linkage hypothesis, where a change from NS A to NS B may have conferred some type of fitness advantage, promoting the genome constellation with the new NS allele to become fixed and resulting in a local sweep. Despite lineage replacement, previously circulating lineages do not go extinct, but rather have a marked decrease in frequency in the system which may be explained by negative frequency dependent selection (Gandon & Michalakis, 2002). That is, the fitness of a phenotype or genotype decreases as it becomes more common, likely due to increased immunity against these lineages in the population. In addition to competitive interactions between lineages, the migratory behaviour of avian hosts and their distinct geographic distribution may contribute to lineage maintenance in different host populations and to the patterns of replacement between lineages in one site. Given these lineages do not go extinct, it is likely they circulate in other dabbling duck populations, including Mallard populations, in other geographic locations, including Asia, or in other less characterized host species, such as diving ducks, geese, or even shorebirds, which are not as intensively monitored in surveillance programs.

Unlike in humans, reassortment of AIV (both inter and intra-subtypic reassortment) is prolific in avian systems (Dugan et al, 2008; Macken et al, 2006; Steel & Lowen, 2014; Wille et al, 2013), and is likely critical in both the long-term (decadale scale) and short-term (within year) arms race of AIV against Mallard immunity. Specifically, it is through reassortment that novel HA or NA lineages are introduced and also the mechanism by which entire genome constellations are formed within the population between years. Indeed, it is the generation of novel genome constellations that drives the emergence of pandemic influenza viruses in human populations (Smith et al, 2009; Wille & Holmes, 2020). In our study, the role of reassortment is most evident when assessing H4N6 genome constellations within a year. Specifically, during periods of high infection burden of H4 viruses in the mallard population, we may expect that the dominant genome constellation has high fitness allowing for rapid proliferation in the mallard population. However, within ~2 weeks of entering/detection in the mallard population the initially dominant H4 constellation rapidly unlinks through reassortment, and is replaced by a variety of auxillary genome constellations. We hypothesize that this is driven by increasing immunity in the host population, resulting in a decrease in fitness for this common constellation. While most antibodies are directed at the HA, and to a lesser extent the NA, during infection all influenza proteins are expressed in infected cells and can potentially induce an antibody response (Krammer, 2019). However, some proteins are more accessible than others, including the NP and M segments (Haaheim, 1977; Sukeno et al, 1979). There is some evidence from monoclonal antibody isolation and from antigenic fingerprinting that natural infection also induces antibodies against internal segment proteins, although the magnitude and quality of these responses is not well defined (Krammer, 2019; Krejnusova et al, 2009; Thathaisong et al, 2008; Yodsheewan et al, 2013). As such, it is likely that through the generation of novel genome constellations, the H4 subtype may be maintained in the population beyond the point at which there is an increase HA-directed immunity and a subsequent decrease in viral fitness. Novel segments incorporated into novel genome constellations originate from other viruses co-circulating in the mallard population (Wille et al, 2013). Indeed, in years with the highest number of mixed infections, a number of other subtypes were co-circulating at relatively high frequency: H1N1 in 2006 and H11N2 in 2008.

### Do genetic patterns reflect antigenic patterns?

Both the phenomenon of lineage replacement and reassortment are important drivers in this system, and would only be selected for if they provided a fitness advantage. One such advantage would be escape from acquired population immunity. Following infection, mallards develop a immune response that is homosubtypic, with potential for heterosubtypic effects (Latorre-Margalef et al, 2017; Latorre-Margalef et al, 2013). Although strength and duration of the immune response is largely unknown in ducks, it is hypothesized that immune memory may be weak and short-lived (Magor, 2011). However, repeated sampling of individual sentinel mallards illustrate maintainance of anti-NP antibodies for months following AIV infection. Furthermore, these same mallards were unlikely to be reinfected with the same subtypes in their second year (Tolf et al, 2013). Indeed, it is hypothesized that this pattern of both homo- and heterosubtypic immunity drives the temporal sucession of different HA lineages across the sampling season (Latorre-Margalef et al, 2014).

Based on the phenomenon of lineage replacement or succession of H4 between years, we hypothesize that the different HA lineages observed must confer homosubtypic immune escape. Molecular and structural comparisons of sequences in each of the three HA lineages found in the mallard population had substitutions in regions of HA1, corresponding to antigentic sites in the homologous H3 peptide (Broecker et al, 2018). Unfortunately, dedicated work on identifying domains has not been completed for H4, however H3 and H4 comprise the H3 Clade of Group 2 viurses (Latorre-Margalef et al, 2013), such that we predict the antigenic sites of H3 should correspond to H4 as well. We found that the 6 to 9 amino acid differences delineating the H4 lineages described in this study partly map to different regions in the crystal structure of H4 HA1, where substitutions in the 2007 sequence were located at the top of the globular head and substitutions in 2008 and 2009 sequences located closer to the stem region of HA1. This may suggest that at a few changes in structural regions in putative antigenic sites is sufficent to evade immunity. Indeed, from work on human influenza A virus H3, it is clear that only a few substitutions are required for immune escape in humans, given they are located in or adjacent to the receptor binding domain (Koel et al, 2013; Smith et al, 2004). In contrast to the HA of human influenza A (*e.g.* (Koel et al, 2013), antigenic diversity of avian HAs has not been well explored, with a few exceptions (*e.g.* (Hill et al, 2016; Koel et al, 2014; Verhagen et al, 2020). However, a recent study investigating infection probability given previous exposure illustrated H3 vaccine escape despite >95% sequence differences in ducks (Wille et al, 2016) In this study, ducks were vaccinated with an inactivated H3 virus, isolated in 2010, and were challenged with naturally circulating AIVs in 2013. Despite a raised immune response due to the vaccination, ducks were not protected against H3 viruses and it was hypotheised that the lack of immunity was associated to substitutions in immunogenic epitopes of more recent H3 strains (e.g. vaccine escape) (Wille et al, 2016) . Despite the parallels, without performing a characterization of H4 antigenic sites or through antigenic cartography, we are unable to confirm whether the different lineages (L1-L3) of H4 confer antigenic differences, and/pr whether reassortment confers partial escape from population immunity.

Understanding of the evolutionary processes of AIV is critical. In humans, the detailed interrogation of seasonal influenza viruses are key to vaccine selection and detection of antiviral resistance (Koel et al, 2013; Van Poelvoorde et al, 2020; Vijaykrishna et al, 2015; Wille & Holmes, 2020). On a larger scale, it is through the process of reassortment that novel pandemic viruses are generated (Gething et al, 1980; Smith et al, 2009). Within the avian reservoir, there is a key focus on HA subtypes that cause significant morbidity and mortality in food production birds (eg H5, H7) or have zoonotic potential (eg H5, H7, H9) (Chang et al, 2020; Chang et al, 2018; Cui et al, 2020; Mahmoud et al, 2019). We generally have a limited understanding of the evolution of LPAIV in the wild bird reservoir (Chen & Holmes, 2006, 2010; Dugan et al, 2008; Wille et al, 2013). But genetic diversity is shaped by virus-related factors including antigenic drift and reassortment, as well as immunological and ecological features of the hosts such as migratory behaviour (van Dijk et al, 2018). Low pathogenicity viruses circulating in wild birds play an important role by providing genetic variation to all influenza A viruses. For example, all human pandemic influenza A viruses have had least one gene of avian origin (Kawaoka et al, 1989; Lindstrom et al, 2004; Rabadan et al, 2006; Runstadler et al, 2013; Scholtissek et al, 1978; Taubenberger et al, 2005), and HPAIV emerge following rapid evolution of LPAIV in poultry (Seekings et al, 2018). Thus, in order to understand evolutionary factors governing AIV dynamics, including host ecology affects, long-term evolution, and genetic structure, it is pivotal to characterize viruses circulating in the wild bird reservoir.

## Acknowledgments

This study was supported by grants from the Swedish Environmental Protection Agency (V-124-01 and V-98-04), the Swedish Research Council VR (2008-58, 2010-3067, 2011-48), the Swedish Research Council Formas (2007-297, 2009-1220) and the Sparbanksstiftelsen Kronan. The surveillance at Ottenby was part of the European Union wild bird surveillance and has received support from the Swedish Board of Agriculture and from the EU FP6-funded New Flubird project. Sequencing has been funded in whole or part with federal funds from the National Institute of Allergy and Infectious Diseases, National Institutes of Health, Department of Health and Human Services under contract number HHSN272200900007C. MW was supported by a PGS-D2 from the Natural Sciences and Engineering Research Council (NSERC) (2011-2013), and is currently supported by an Australian Research Council Discovery Early Career Research Award (DE200100977). OGP and JR were supported by the GCRF One Health Poultry Hub (Grant No. BB/S011269/1).This manuscript was prepared while DEW was employed at the JCVI. The opinions expressed in this article are the author’s own and do not reflect the view of the Centers for Disease Control, the Department of Health and Human Services, or the United States government

We thank all the staff at Ottenby Bird Observatory for trapping and sampling Mallards, and all laboratory staff for processing samples over the years.

This is manuscript 3XX from Ottenby Bird Observatory.

